# Native American Ancestry and Pigmentation Allele Contributions to Skin Color in a Caribbean Population

**DOI:** 10.1101/2021.11.24.469305

**Authors:** Khai C Ang, Victor A Canfield, Tiffany C Foster, Thaddeus D Harbaugh, Kathryn A Early, Rachel L Harter, Katherine P Reid, Shou Ling Leong, Yuka I Kawasawa, Dajiang J Liu, John W Hawley, Keith C Cheng

**Affiliations:** Department of Pathology, Penn State College of Medicine, Hershey, PA, USA; Jake Gittlen Laboratories for Cancer Research, Penn State College of Medicine, Hershey, PA, USA; Department of Family & Community Medicine, Penn State College of Medicine, Hershey, PA, USA; Department of Biochemistry and Molecular Biology, Penn State College of Medicine, Hershey, PA, USA; Department of Pharmacology, Penn State College of Medicine, Hershey, PA, USA; Institute of Personalized Medicine, Penn State College of Medicine, Hershey, PA, USA; Department of Public Health Sciences, Penn State College of Medicine, Hershey, PA, USA; Salybia Mission Project, Dominica

## Abstract

Our interest in the genetic basis of skin color variation between populations led us to seek a Native American population with African admixture but low frequency of European light skin alleles. Analysis of 458 genomes from individuals residing in the Kalinago territory of the Commonwealth of Dominica showed approximately 55% Native American, 32% African, and 12% European ancestry, the highest Native American ancestry among Caribbean populations to date. Skin pigmentation ranged from 20 to 80 melanin units, averaging 46. Three albino individuals were determined to be homozygous for a causative multi-nucleotide polymorphism *OCA2*^*NW273KV*^ contained within a haplotype of African origin; its allele frequency was 0.03 and single allele effect size was -8 melanin units. Derived allele frequencies of *SLC24A5*^*A111T*^ and *SLC45A2*^*L374F*^ were 0.14 and 0.06, with single allele effect sizes of -6 and -4, respectively. Native American ancestry by itself reduced pigmentation by more than 20 melanin units (range 24 - 29). The responsible hypopigmenting genetic variants remain to be identified, since none of the published polymorphisms predicted in prior literature to affect skin color in Native Americans caused detectable hypopigmentation in the Kalinago.

## Introduction

Human skin pigmentation is a polygenic trait that is influenced by health and environment (Barsh, 2003). Lighter skin is most common in populations adapted to northern latitudes characterized by lower UV incidence than equatorial latitudes (Jablonski and Chaplin, 2000). Selection for lighter skin, biochemically driven by a solar UV-dependent photoactivation step in the formation of vitamin D (Engelsen, 2010; Hanel and Carlberg, 2020; Holick, 1981; Loomis, 1967) is regarded as the most likely basis for a convergent evolution of lighter skin color in European and East Asian/Native American populations (Lamason et al., 2005; Norton et al., 2007). The hypopigmentation polymorphisms of greatest significance in Europeans have two key characteristics: large effect size and near fixation. For example, the *A111T* allele in *SLC24A5* (Lamason et al., 2005) explains at least 25% of the difference in skin color between people of African vs. European ancestry, and is nearly fixed in European populations. No equivalent polymorphism in Native Americans or East Asians has been found to date.

Native Americans share common ancestry with East Asians (Derenko et al., 2010; Tamm et al., 2007), diverging before ∼15 kya (Gravel et al., 2013; Moreno-Mayar et al., 2018; Reich et al., 2012), but the extent to which these populations share pigmentation variants remains to be determined. The derived alleles of rs2333857 and rs6917661 near *OPRM1*, and rs12668421 and rs11238349, in *EGFR* are near fixation in some Native American populations, but all also have a high frequency in Europeans (Quillen et al., 2012), and none reach genome-wide significance in Adhikari et al., (2019). However, the latter found a significant association for the *Y182H* variant of *MSFD12* with skin color, but its frequencies were only 0.27 and 0.17 in Native Americans and East Asians, respectively, suggesting that it can explain only a small portion of the difference between Native American and/or East Asians and African skin color. Thus, the genetic basis for lighter skin pigmentation specific to Native American and East Asian populations, whose African alleles would be expected to be ancestral, remains to be found.

The shared ancestry of East Asians and Native Americans suggests the likelihood that some light skin color alleles are shared between these populations. This is particularly the case for any variants that achieved fixation in their common ancestors. For Native American populations migrating from Beringia to the Tropics, selection for darker skin color also appears likely (Jablonski and Chaplin, 2000; Quillen et al., 2018). This would have increased the frequency of novel dark skin variants, if any, and would have decreased the frequency of light skin variants that had not achieved fixation. Hypopigmenting alleles are associated with the European admixture characteristic of many current Native American populations (Brown et al., 2017; Gravel et al., 2013; Keith et al., 2021; Klimentidis et al., 2009; Reich et al., 2012). Since the European hypopigmenting alleles may mask the effects of East Asian and Native American alleles, we searched for an admixed Native American population with high African, but low European admixture.

Prior to European contact, the Caribbean islands were inhabited by populations who migrated from the northern coast of South America (Benn-Torres et al., 2008; Harvey et al., 1969; Honychurch, 2012; “Island Caribs,” 2016; Torres et al., 2015, 2013). During the Colonial period, large numbers of Africans were introduced into the Caribbean as slave labor (Honychurch, 2012; Torres et al., 2013). As a consequence of African and European admixture and high mortality among the indigenous populations, Native American ancestry now contributes only a minor portion (<15%) of the ancestry of most Caribbean islanders (The 1000 Genomes Project Consortium, 2015, 2010; Torres et al., 2015, 2013). The islands of Dominica and St. Vincent were the last colonized by Europeans in the late 1700s (Honychurch, 2012, 1998; Rogoziński, 2000). In 1903, the British granted 15 km^2^ (3,700 acres) on the eastern coast of Dominica as a reservation for the Kalinago, who were then called “Carib”. When Dominica gained Independence in 1978, legal rights and a degree of protection from assimilation were gained by the inhabitants of the Carib Reserve (Honychurch, 2012) (redesignated *Kalinago Territory* in 2015). Oral history and beliefs among the Kalinago, numbering about 3,000 living within the Territory (“Kalinago Territory,” 2021) (Figure S1) are consistent with the primarily Native American and African ancestry, assessed and confirmed genetically here.

Early in our genetic and phenotypic survey of the Kalinago, we noted an albino individual, and upon further investigation, we learned of two others residing in the Territory. We set out to identify the mutant albinism allele to avoid single albino allele effects that would potentially mask Native American hypopigmentation alleles. Oculocutaneous albinism (OCA) is a recessive trait characterized by visual system abnormalities and hypopigmentation of skin, hair, and eyes (Gargiulo et al., 2011; Gronskov et al., 2007; Grønskov et al., 2014; Hong et al., 2006; Vogel et al., 2008) that is caused by mutations in any of a number of autosomal pigmentation genes (Carrasco et al., 2009; Edwards et al., 2010; Gao et al., 2017; Grønskov et al., 2013; Kausar et al., 2013; King et al., 2003; Spritz et al., 1995; Stevens et al., 1997, 1995; Vogel et al., 2008; Woolf, 2005; Yi et al., 2003). The Incidence of albinism is ∼1:20,000 in populations of European descent, but much higher in some populations, including many in sub-Saharan Africa (1:5,000)(Greaves, 2014). Here, we report on the ancestry of a population sample representing 15% of the Kalinago population of Dominica, the identification of the new albinism allele in that population, and measurement of the hypopigmenting effects of the responsible albinism allele, the European *SLC24A5*^*A111T*^ and *SLC45A2*^*L374*^ alleles. Native American ancestry alone caused a measurable effect on pigmentation. In contrast, alleles identified in past studies of Native American skin color caused no significant effect on skin color.

## Results & Discussion

Our search for a population admixed for Native American/African ancestries with minimal European admixture led us to the “Carib” population in the Commonwealth of Dominica. Observations from an initial trip to Dominica suggested wide variation in Kalinago skin color. Pursuit of the genetic studies described here required learning about oral and written histories, detailed discussion with community leadership, IRB approval from Ross University (until Hurricane Maria in 2017, the largest medical school in Dominica) and the Department of Health of the Commonwealth of Dominica, and relationship-building with three administrations of the Kalinago Council over 15 years.

### Population Sample

Our DNA and skin-color sampling program encompassed 458 individuals, representing 15% of the population of the territory and all three known albino individuals. Ages ranged from 6 to 93 (Table S1 and Figure S2). We were able to obtain genealogical information for about half of the parents (243 mothers and 194 fathers). Community-defined ancestry (described as ‘Black,’ ‘Kalinago,’ or ‘Mixed’) for both parents was obtained for 426 individuals (92% of sample), including 108 parents from whom DNA samples were obtained (72 Kalinago, 36 Mixed, and 0 Black).

### Kalinago Ancestry

The earliest western mention of the Kalinago (originally as “Caribs”) was in Christopher Columbus’s journal dated 26th November 1492 (Honychurch, 2012). Little is known about the detailed cultural and genetic similarities and differences between them and other Caribbean pre-contact groups such as the Taino. African admixture in the present Kalinago population derived from the African slave trade; despite inquiry across community, governmental, and historical sources, we were unable to find documentation of specific regions of origin in Africa or well-defined contributions from other groups. The population’s linguistics are uninformative, as they speak, in addition to English, the same French-based Antillean Creole spoken on the neighboring islands of Guadeloupe and Martinique.

To study Kalinago population structure, we analyzed an aggregate of our Kalinago SNP genotype data and HGDP data (Li et al., 2008) using ADMIXTURE (Figures 1 and S3) as described in Methods. At K=3, the ADMIXTURE result confirmed the three major clusters, corresponding roughly to Africans (blac cluster), European/ Middle Easterners/ Central & South Asians (yellow cluster), and East Asians/ Native Americans (green cluster). At K=4 and higher, the Native American component that predominates in Kalinago (red cluster) separates from the East Asians (green cluster). Consistent with prior work (Li et al., 2008), an Oceanian component (purple cluster) appears at K=5 and a Central & South Asian component (brown cluster) appears at K=6; both are minor sources of ancestry in our Kalinago sample (average <1%) (Table S2).

**Figure 1:**
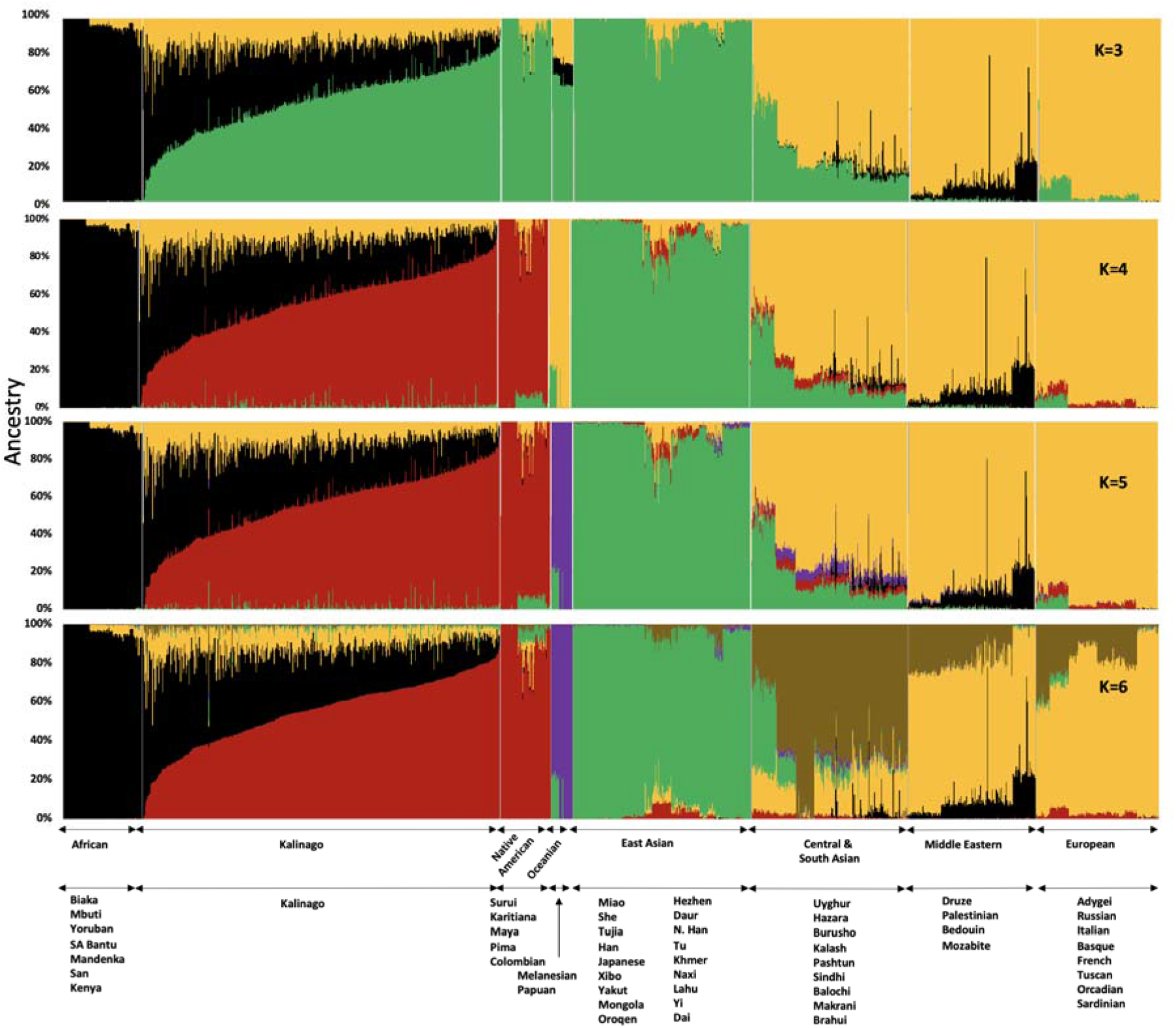
Admixture analysis of Kalinago compared with Human Genome Diversity Project populations. Results are depicted using stacked bar plots, with one column per individual. At K=3, the Kalinago, Native Americans, Oceanians, and East Asians fall into the same green cluster. At K=4, the Native Americans (red cluster) are separated from the East Asians (green cluster).

At K=4 to K=6, the Kalinago show on average 55% Native American, 32% African, and 11-12% European ancestry. Estimates from a two-stage admixture analysis are similar, as are results from local ancestry analysis (see Methods) (Table S3), leading to estimates of 54-56% Native American, 31-33% African, and 11-13% European ancestry. The individual with the least admixture has approximately 94% Native American and 6% African ancestry. The results of the principal component (PC) analysis (Figure S4) were consistent with ADMIXTURE analysis. The first two PCs suggest that most Kalinago individuals show admixture between Native American and African ancestry, with a smaller but highly variable European contribution apparent in the displacement in PC2 (Figure S4A). A smaller number of Kalinago individuals with substantial East Asian ancestry exhibit displacement in PC3 (Figure S4B).

Our analysis of Kalinago ancestry revealed considerably more Native American and less European ancestry than the Caribbean samples of Torres et al. (2013) and the admixed populations from the 1000 Genomes Project (The 1000 Genomes Project Consortium, 2015) (Figure 2). Some Western Hemisphere Native Americans reported in Reich et al. (2012) have varying proportions of European but very little African admixture (Figure 2B). Overall, the Kalinago have more Native American and less European ancestry than any other Caribbean population.

**Figure 2.**
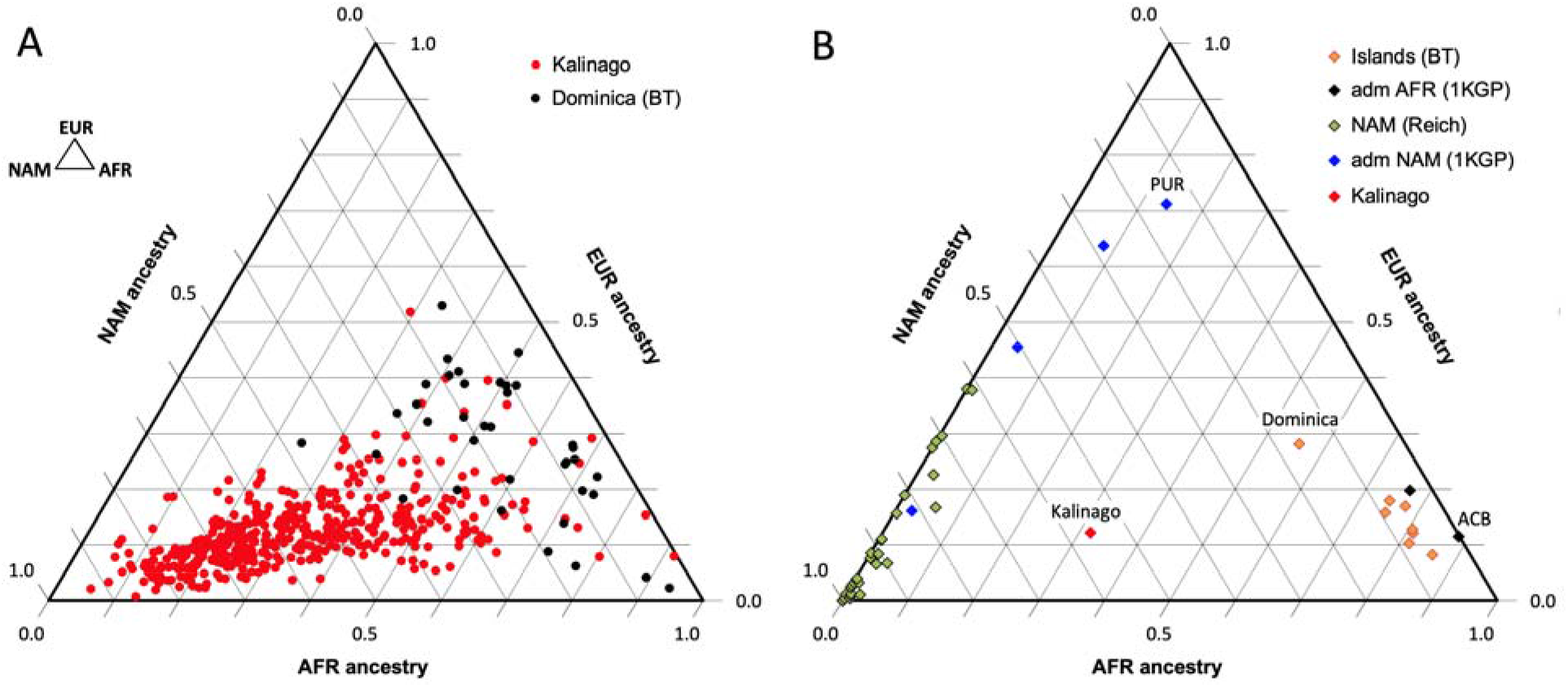
Comparison of Kalinago ancestry with that of other populations in the Western Hemisphere. Ternary plots of ancestry from our work and the literature show estimated proportions of African (AFR), European (EUR) and Native American (NAM) ancestry. *A*, Comparison of individuals (n=452, omitting 6 individuals with EAS > 0.1) genotyped in this study to individuals (n=38) from southern Dominica sampled by Torres et al., 2013, *B*, Comparison of the Kalinago average ancestry with other Native American populations. Kalinago, this study (n=458); Islands (BT) indicates Caribbean islanders reported in Torres et al., 2013, with Dominica labeled; admixed (adm) AFR (1KGP) and admixed NAM (1KGP) represent admixed populations from The 1000 Genomes Project Consortium, 2015, with Caribbean samples PUR (Puerto Rico) and ACB (Barbados) labeled; and AMR (Reich) indicates mainland Native American samples reported in Reich et al., 2012. Inset (top left) shows ancestries at vertices.

The 55% Native American ancestry calculated from autosomal genotype in the Kalinago is greater than the reported 13% in Puerto Rico (Gravel et al., 2013), 10-15% for Tainos across the Caribbean (Schroeder et al., 2018), and 8% for Cubans (Marcheco-Teruel et al., 2014). This is also considerably higher than the reported 6% Native American ancestry found in Bwa Mawego, a horticultural population that resides south of the Kalinago Territory (Keith et al., 2021). However, this result is lower than the 67% Native American ancestry reported by Crawford et al. (2021) for an independently collected Kalinago samples based on the mtDNA haplotype analysis. This difference suggests a paternal bias in combined European and/or African admixture. Since our Illumina SNP-chip genotyping does not yield reliable identification of mtDNA haplotypes, we are currently unable to compare maternal to autosomal ancestry proportions for our sample. Samples genotyped using 105 ancestry informative markers from Jamaica and the Lesser Antilles (Torres et al., 2015, 2013) yielded an average of 7.7% Native American ancestry (range 5.6% to 16.2%), with the highest value from a population in Dominica sampled outside the Kalinago reservation. Relevant to the potential mapping of Native American light skin color alleles, the Kalinago population has among the lowest European ancestry (12%) compared to other reported Caribbean Native Americans in St. Kitts (8.2%), Barbados (11.5%) and Puerto Rico (71%) (Torres et al., 2013). Contributing to the high percentage of Native American ancestry in the Kalinago is their segregation within the 3,700-acre Kalinago Territory in Dominica granted by the British in 1903, and the Kalinago tradition that women marrying non-Kalinago are required to leave the Territory; non-Kalinago spouses of Kalinago men are allowed to move to the Territory (KCA, KCC, Personal Communication with Kalinago Council, 2014). These factors help to explain why samples collected outside the Kalinago territory (Torres et al., 2013), show lower fractional Native American ancestry.

During our fieldwork, it was noted that members of the Kalinago community characterized themselves and others in terms of perceived ancestry perceived as “Black,” “Kalinago,” or “Mixed.” Compared to individuals self-identified as “Mixed,” those self-identified as “Kalinago” have on average more Native American ancestry (67% vs 51%), less European ancestry (10% vs 14%), and less African ancestry (23% vs 34%) (Figure S5). Thus, these folk categories based on phenotype are reflected in some underlying differences in ancestry (genotype).

### Kalinago Skin Color Variation

Melanin index unit (MI) calculated from skin reflectance measured at the inner upper arm (see Methods) was used as a quantitative measure of melanin pigmentation (Ang et al., 2012; Diffey et al., 1984). MI determined in this way Is commonly used as a measure of constitutive skin pigmentation (Choe et al., 2006; Park and Lee, 2005). The MI in the Kalinago ranged from 20.7 to 79.7 (Figure S6), averaging 45.7. The three Kalinago albino individuals sampled had the lowest values (20.7, 22.4 and 23.8). Excluding these, the MI ranged between 28.7 to 79.7 and averaged 45.9. For comparison, the MI averaged 25 and 21 for people of East Asian and European ancestry, respectively, as measured with the same equipment in our laboratory (Ang et al., 2012; Tsetskhladze et al., 2012). This range is similar to that of another indigenous population, the Senoi of Peninsular Malaysia (MI 24 to 78; mean = 45.7) (Ang et al., 2012). The Senoi are believed to include admixture from Malaysian Negritos whose pigmentation is darker (mean = 55) (Ang et al., 2012) than that of the average Kalinago. In comparison, the average MI was 53.4 for Africans in Cape Verde (Beleza et al., 2012) and 59 for African-Americans (Shriver et al., 2003). Individuals described as “Kalinago” were slightly lighter and had a narrower MI distribution (42.5 ± 5.6, mean ± SD) compared to “Mixed” (45.8 ± 9.6) (Figure S7).

### An OCA2 albinism allele in the Kalinago

Oculocutaneous albinism (OCA) is a genetically determined condition characterized by nystagmus, reduced visual acuity, foveal hypoplasia and strabismus as well as hypopigmentation of the skin, hair and eye (Dessinioti et al., 2009; van Geel et al., 2013). The three sampled albino individuals had pale skin (MI 20.7, 22.4 and 23.8 vs. 29-80 for non-albino individuals), showed nystagmus, and reported photophobia and high susceptibility to sunburn. In contrast to the brown irides and black hair of most Kalinago, including their parents, the albino individuals had blonde hair and grey irides with varying amounts of green and blue.

To identify the albinism variant in the Kalinago, we first determined that none of the albino individuals carried any of 28 mutations previously found in African or Native American albino individuals (Carrasco et al., 2009; King et al., 2003; Stevens et al., 1997; Yi et al., 2003), including a 2.7 kb exon 7 deletion in *OCA2* found at high frequency in some African populations. Whole exome sequencing of one albino individual and one parent (obligate carrier) revealed polymorphisms homozygous in the albino individuals and heterozygous in the parent, an initial approach that assumes that the albino individual was not a compound heterozygote. We identified 12 variant alleles in 7 oculocutaneous albinism genes (or genomic regions) that met these criteria (summarized in Table S4A). None were nonsense or splice site variants. Five of the twelve variants were intronic, one was synonymous, one was located in 5’UTR, and three were in the 3’UTR (Table S4B). Two missense variants were found in *OCA2*: SNP rs1800401 (c.913C>T or p.Arg305Trp in exon 9), *R305W*, and multi-nucleotide polymorphism rs797044784 in exon 8 (c.819_822delCTGGinsGGTC; p.Asn273_Trp274delinsLysVal), *NW273KV*.

Among 458 Kalinago *OCA2* genotypes, 26 carried *NW273KV* and 60 carried *R305W* (Table 1). Only *NW273KV* homozygotes were albino individual. We know that the allele responsible for albinism was *NW273KV* because neither of the two individuals homozygous for *R305W* but not *NW273KV*, was albino individual. In further support of this conclusion is that one individual who was homozygous for *R305W* and homozygous ancestral for *NW273* had an MI of 72, among the darkest in the entire population. *R305W* is notably present with frequency > 0.10 in some African, South Asian, and European populations (The 1000 Genomes Project Consortium, 2015), predicting a Hardy-Weinberg frequency of homozygotes above 1%. This is far greater than the observed frequency of individuals with albinism and therefore inconsistent with the idea that this is a deleterious variant. The fact that *R305W* scores incorrectly as pathogenic using SIFT, Polyphen 2.0 and PANTHER that *R305W* (Kamaraj and Purohit, 2013) indicates a need for refinement of these methods. The universal association of *R305W* with the *NW273KV* haplotype indicates that the founder haplotype of the *NW273KV* albinism mutation carried the silent *R305W* variant.

**Table 1.** Albinism among NW273KV and R305W genotypes.

| Allele/Genotype |  | ***NW273KV* genotype** |  |  | Total |
| --- | --- | --- | --- | --- | --- |
|  |  | Homozygous Ancestral<sup>a</sup> | Heterozygous | Homozygous Derived |  |
| ***R305W* genotype** | Homozygous Ancestral | 398 | 0 | 0 | 398 |
|  | Heterozygous | 33 | 22 | 0 | 55 |
|  | Homozygous Derived | 1 | 1 | 3* | 5 |
|  | Total | 432 | 23 | 3* | 458 |

<sup>a</sup> Ancestral=reference allele and derived=alternate allele for both variants.
\* Albino phenotype. Notably, none of the other genotypic categories are albino individuals

To identify the origin of the albino allele, albino individuals and carriers were analyzed for regions exhibiting homozygosity, and identity-by-descent and local ancestry was estimated (see Methods). All three albino individuals share a homozygous segment of ∼1.7 Mb that encompasses several genes in addition to OCA2 (Figure 3). The albino haplotype, defined by homozygosity in individuals 2 and 3 extends ∼11 Mb; comparison to local ancestry shows that this haplotype is clearly of African origin.

**Figure 3.**
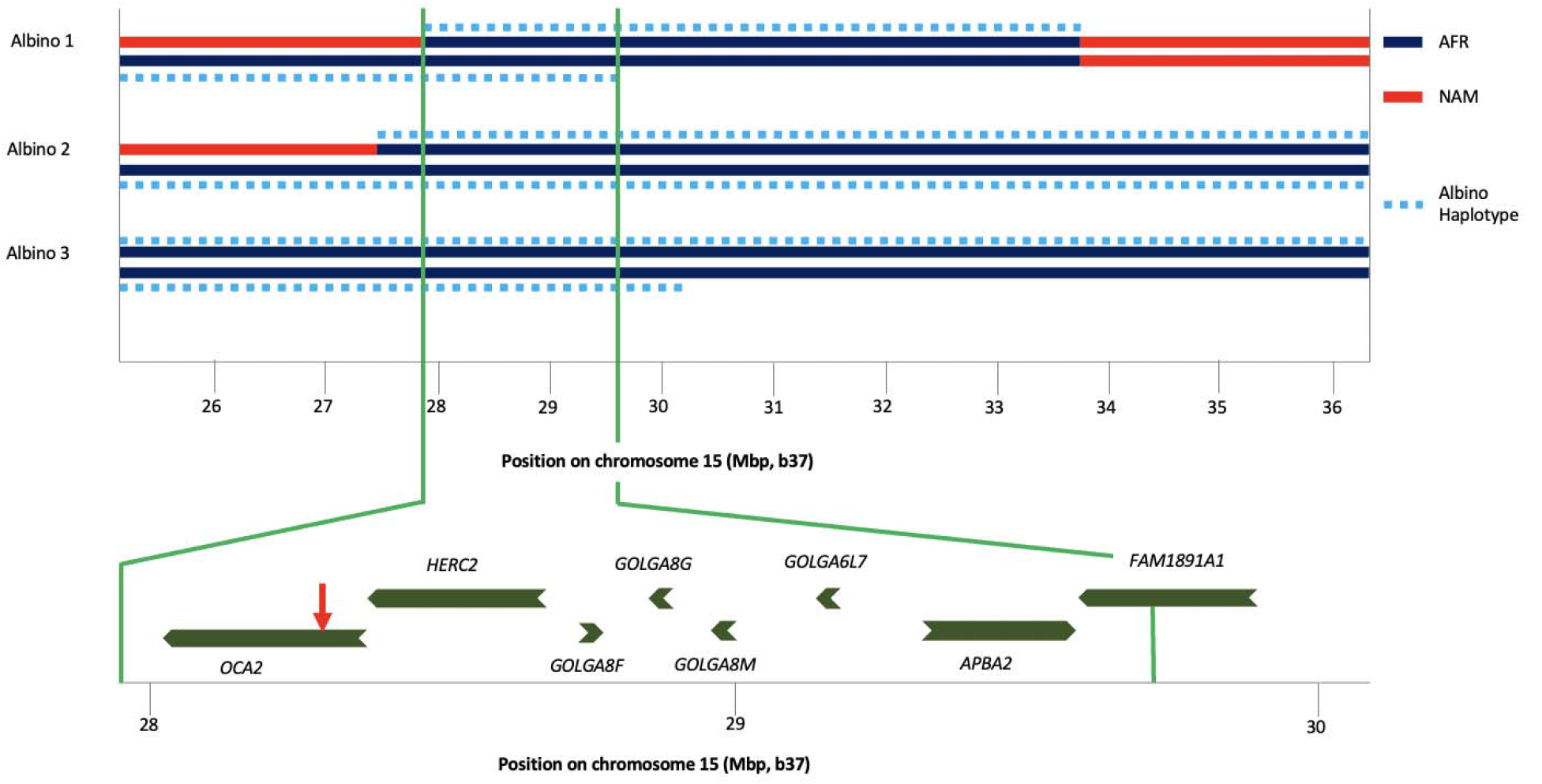
Haplotype analysis for three albino individuals. The inner two lines indicate NAM (red) or AFR (dark blue) ancestry; no EUR ancestry was found in this genomic region. For this local ancestry analysis, the region shown here consisted of 110 non-overlapping segments with 7 to 346 SNPs each (mean 65). The deduced extent of shared albino haplotype (dotted light blue lines) is indicated on each chromosome. The common region of overlap indicated by the minimum homozygous region (determined by albino individual 1) shared by all three albino individuals is shown at expanded scale below. Genes in this region are labeled, and the position of the NW273KV polymorphism in OCA2 is indicated by the red arrowhead.

The Kalinago albino individuals are the only reported individuals where the albinism was caused by homozygosity for the NW273KV allele of OCA2. Two reported albino individuals of African-American/Dutch descent were compound heterozygotes for the OCA2 mutation, with one allele being the NW273KV variant chromosome (Garrison et al., 2004; Lee et al., 1994). Conservation of the NW sequence among vertebrates and its inclusion in a potential N-linked glycosylation site (Rinchik et al., 1993) that is eliminated by the mutation supports the variant’s pathogenicity. The NW273KV frequency in our sample (0.03) translates into a Hardy-Weinberg albinism frequency (p^2^ = 0.0009) of ∼1 per 1000, as observed (3 in a population of about 3000). Examination of publicly available data reveals three *OCA2*^*NW273KV*^ heterozygotes in the 1000 Genome Project, a pair of siblings from Barbados (ACB) and one individual from Sierra Leone (MSL) (The 1000 Genomes Project Consortium, 2015). The three 1KGP individuals share a haplotype of ∼1.5 Mb, of which ∼1.0 Mb matches the albino haplotype in the Kalinago. The phasing for the *OCA2*^*NW273KV*^ variant in the public data is inconsistent, with the variant assigned to the wrong chromosome for the ACB siblings.

### Genetic Contributions to Kalinago Skin Color Variation

One motivation for undertaking this work was to characterize genetic contributions to skin pigmentation in a population with primarily Native American and African ancestry, so that we could focus on the effect of Native American hypopigmenting alleles without interference from European alleles. The Kalinago population described here comprises the only population we are aware of that fits this ancestry profile. To control for the effects of the major European pigmentation loci, all Kalinago samples were genotyped for *SLC24A5*^*A111T*^ and *SLC45A2*^*L374F*^. The phenotypic effects of these variants and *OCA2*^*NW273KV*^ are shown in Figure 4. Each variant decreases melanin pigmentation, with homozygotes being lighter than heterozygotes. The greatest effect is seen in the *OCA2*^*NW273KV*^ homozygotes (the albino individuals), as previously noted. The frequencies of the derived alleles of *SLC24A5*^*A111T*^ and *SLC45A2*^*L374F*^ in the Kalinago sample are 0.14 and 0.06, respectively.

**Figure 4.**
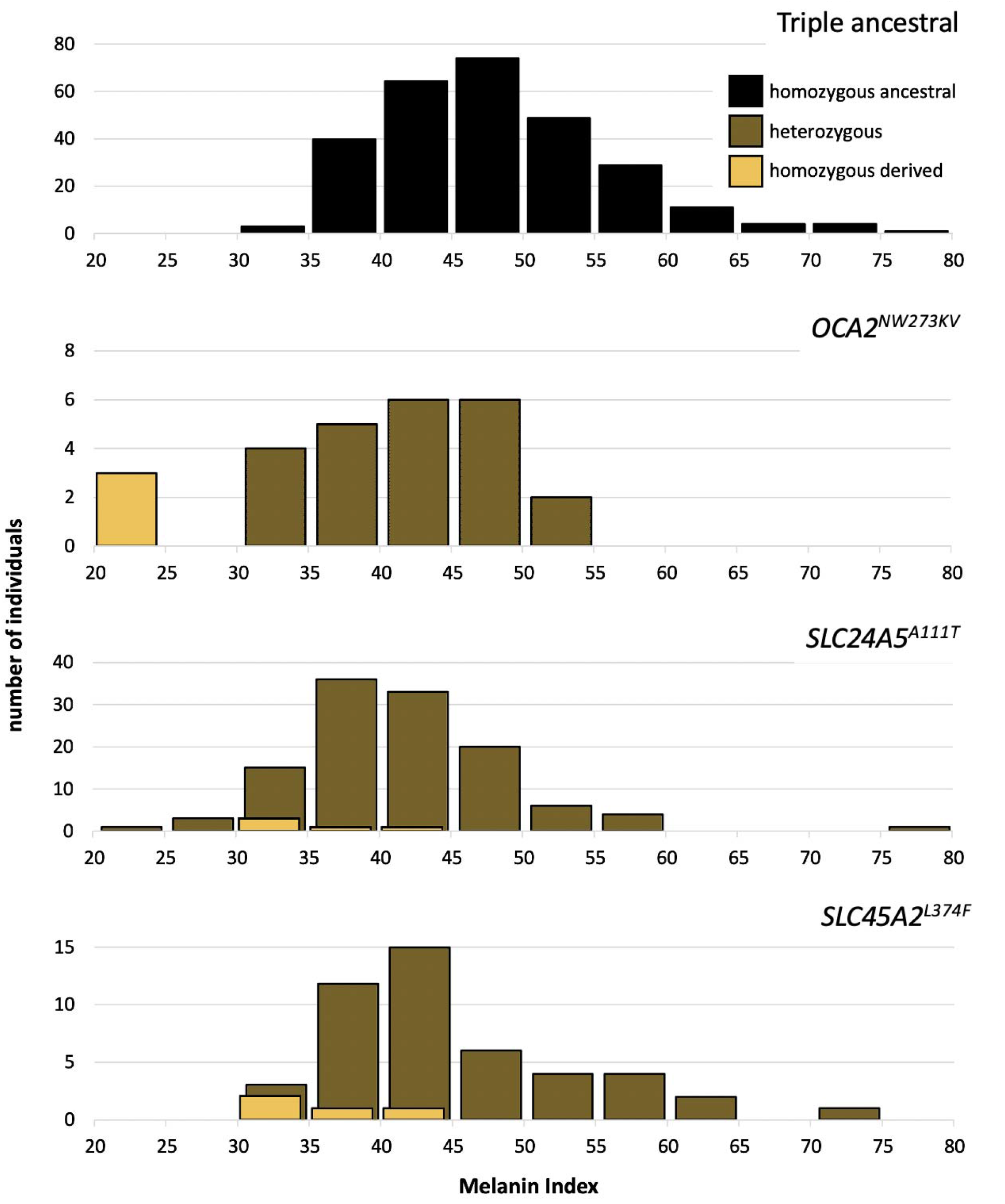
Skin color distribution of Kalinago samples according to genotype. The “Triple ancestral” plot is individuals ancestral for three pigmentation loci (SLC24A5^111A^, SLC45A2^374L^ and OCA2^273NW^). In the other plots, heterozygosity or homozygosity is indicated for the variants: OCA2^NW273KV^; SLC24A5^A111T^; and SLC45A2^L374F^. Individuals depicted in the 2^nd^ through 4^th^ panels are repeated if they carry variants at more than one locus.

The markedly higher frequency of *SLC24A5*^*A111T*^ compared to *SLC45A2*^*L374F*^ is not explained solely by European admixture, given that most Europeans are nearly fixed for both alleles. This deviation can be explained by the involvement of source populations that carry the *SLC24A5*^*A111T*^ variant but not *SLC45A2*^*L374F*^. Although some sub-Saharan West African populations (the likeliest source of AFR ancestry in the Kalinago) have negligible *SLC24A5*^*A111T*^ frequencies, moderate frequencies are found in the Mende of Sierra Leone (MSL, allele frequency=0.08) (Micheletti et al., 2020; The 1000 Genomes Project Consortium, 2015), while some West African populations such as Hausa and Mandinka who have allele frequencies of 0.11 and 0.15, respectively (Cheung et al., 2000; Rajeevan et al., 2012). Such African individuals carrying the *SLC24A5*^*A111T*^ allele could potentially cause the observed frequencies by founder effect. In addition, the region of chromosome 5 containing *SLC45A2* exhibits low European ancestry (6.5%) that is consistent with low observed *SLC45A2*^*L374F*^ frequency.

In order to investigate the potential effect of the *SLC25A5*^*A111T*^ allele on the albinism phenotype, we also compared other pigmentation phenotypes such as the hair and eye colors for all albino individuals and carriers. One of the three Kalinago albino individuals was also heterozygous for *SLC24A5*^*A111T*^, but neither skin nor hair color for this individual was lighter than that of the other two albino individuals, who were homozygous for the ancestral allele at *SLC24A5*^*A111*^; this observation is consistent with epistasis of *OCA2* hypopigmentation over that of *SLC24A5*^*A111T*^. Nine sampled non-albino individuals had combinations of hair that was reddish, yellowish, or blonde (n=6), skin with MI < 30 (n=3), and gray, blue, green or hazel irides (n=2); among these, six were heterozygous and one homozygous for *SLC24*A*5*^*A111T*^, and three were heterozygous for the albino variant. A precise understanding of the phenotypic effects of the combinations of these and other hypopigmenting alleles will require further study.

The strong dependence of pigmentation on Native American ancestry is clarified by focusing on individuals lacking the hypopigmenting alleles *SLC24A5*^*A111T*^, *SLC45A2*^*L374F*^, and *OCA2*^*NW273KV*^ (Figure 5). Although positive deviations from the best fit are apparent at both high and low Native American ancestry, the trend toward lighter pigmentation as Native American ancestry increases is clear. The net difference between African and Native American contributions to pigmentation appears likely to be bounded by the magnitudes of the slope vs NAM ancestry (24 units) and the slope vs AFR ancestry (29 units, not shown). The difference in melanin index value is expected to be explained by genetic variants that are highly differentiated between African and Native American populations.

**Figure 5.**
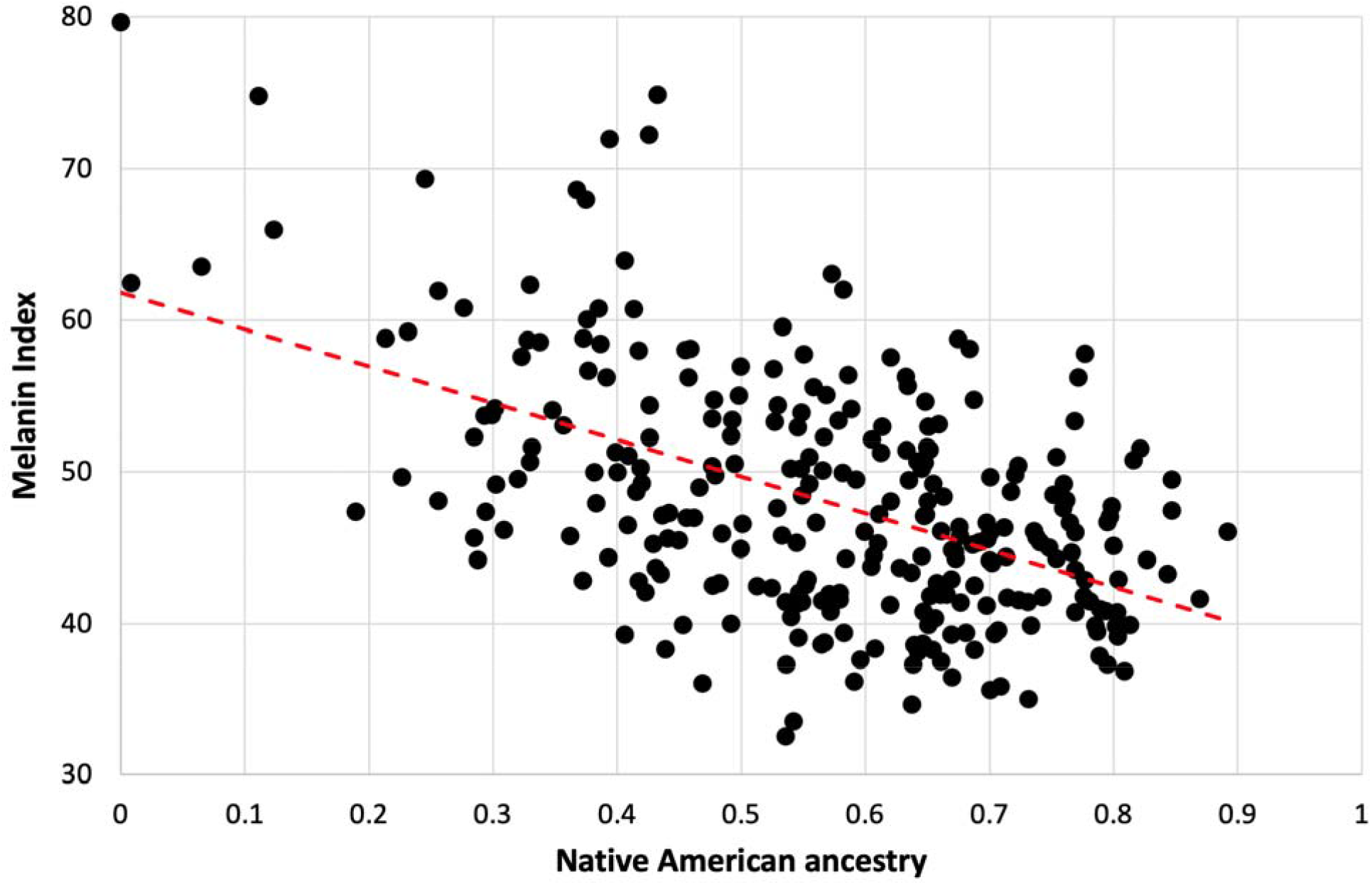
Dependence of Melanin Unit on ancestry for Kalinago. Only individuals who are ancestral for *SLC24A5*^*111A*^, *SLC45A2*^*374L*^, and *OCA2*^*273NW*^ alleles are shown (n=279). The dotted red line represents the best fit (linear regression). Slope is -24.3 (MI = -24.3*NAM + 61.9); r_2_ = 0.2722.

To further investigate the contributions of genetic variation to skin color, we performed association analyses using an additive model for Melanin Index, conditioning on sex, ancestry (using 10 PCs), and genotypes for *SLC24A5*^*A111T*^, *SLC45A2*^*L374F*^ and *OCA2*^*NW273KV*^. Assuming likely epistasis of albinism alleles over other hypopigmenting alleles, these analyses omitted the three albino individuals. Employing a linear regression model, we found that sex and all three genotyped polymorphisms were statistically significant (Table 2 & S5). However, only *SLC24A5*^*A111T*^ reaches genome-wide significance. PC1, which strongly correlated with Native American vs African ancestry, exhibits the lowest *p*-value. Effect sizes were about -6 units (per allele) for *SLC24A5*^*A111T*^, -4 units for *SLC45A2*^*L374F*^ and -8 units for the first *OCA2*^*NW273KV*^ allele.

**Table 2.**
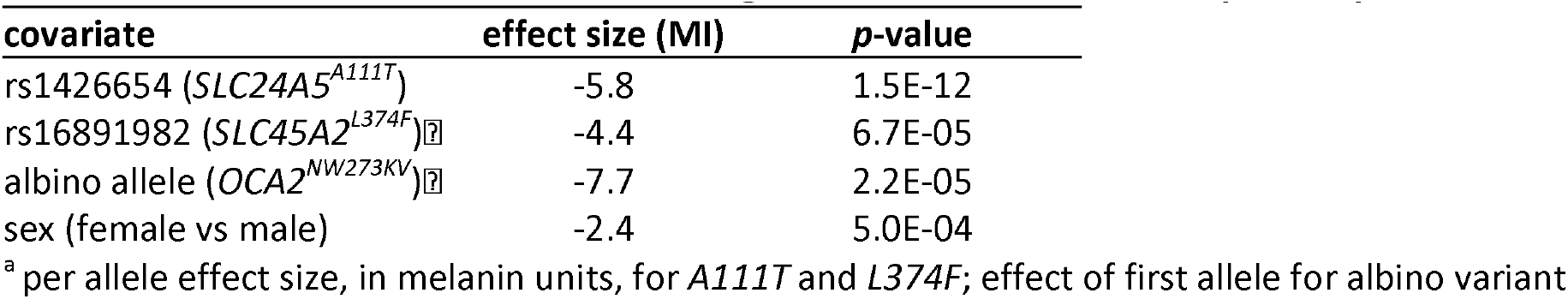
Effect sizes for covariates in linear regression model with 10 Principal Components.

| covariate | effect size (MI) | p-value |
| --- | --- | --- |
| rs1426654 (*SLC24A5*<sup>A111T</sup>) | -5.8 | 1.5E-12 |
| rs16891982 (*SLC45A2*<sup>L374F</sup>)<sup>Ⓜ</sup> | -4.4 | 6.7E-05 |
| albino allele (*OCA2*<sup>NW273KV</sup>)<sup>Ⓜ</sup> | -7.7 | 2.2E-05 |
| sex (female vs male) | -2.4 | 5.0E-04 |

<sup>a</sup> per allele effect size, in melanin units, for *A111T* and *L374F*; effect of first allele for albino variant

Additional covariates were considered but not included in our standard model. Skin pigmentation exhibited a decreasing trend with age, but its contribution was not statistically significant (adjusted P value = 0.08). Estimated effect sizes for significant covariates were little affected by the inclusion of age as a covariate (Table S5). Analysis of SNPs that were previously reported as relevant to pigmentation are shown in Table S6. The lowest (adjusted) *p*-value for this collection of variants is about 0.001, considerably larger than the P values for the variants included as covariates in our standard model. Inclusion of the SNP of lowest *p-*value from each of the five regions containing *BCN2, TYR, OCA2, MC1R*, and *OPRM1* only modestly altered effect sizes for the other covariates (Table S5).

The effect size for *SLC24A5*^*A111T*^ measured here is consistent with previously reported results of -5 melanin units calculated from an African-American sample (Lamason et al., 2005; Norton et al., 2007) and -5.5 from admixed inhabitants of the Cape Verde islands (Beleza et al., 2013). Reported effect sizes for continental Africans are both higher and lower, -7.7 in Crawford et al. (2017) and -3.6 Martin et al. (2017b), while the estimated effect size in the CANDELA study (GWAS of combined admixed populations from Mexico, Brazil, Columbia, Chile and Peru) (Adhikari et al., 2019) yielded an effect size about -3 melanin units.

A significant effect of *SLC45A2*^*L374F*^ on skin pigmentation reported for the African American sample by Norton et al., (2007)and in the CANDELA study by Adhikari et al. (2019) but not for the African Caribbean sample by Norton et al. (2007). The 4 unit effect size of this allele in the Kalinago reported here is similar to the 5 unit effect reported by Norton et al., 2007. Beleza et a. (2013) reported significance for a SNP in strong linkage disequilibrium with *SLC45A2*^*L374F*^, which was itself not genotyped.

Our estimate that a single *OCA2*^*NW273KV*^ allele causes about -8 melanin units of skin lightening is the first reported population-based effect size measurement for any albinism allele. Although albinism is generally considered recessive, our population sample offered an opportunity to compare the effect size for the first and second alleles quantitatively. We applied the estimated parameters to the three albino individuals and found that they were lighter by an average of 10 units than predicted by the additive model (*p* < 0.0024, 1-tailed t-test). The large difference between -10 and -16 means that the additive model for the skin color effect of the albino mutation found in the Kalinago is rejected; the mechanism of this non-linearity and epistasis remains to be fully understood. As expected from the lack of albinism in *R305W* homozygotes, when controlling for *OCA2*^*NW273KV*^ status, *OCA2*^*R305W*^ had no detectable effect on skin color (Table S6).

To identify novel SNPs that may contribute towards skin pigmentation in the Kalinago samples, we performed GWAS using linear regression and linear mixed models (LMMs). Estimated power for these analyses is shown in Figure S8, and Q-Q plots are depicted in Figure S9. The LMM approaches exhibited less statistic inflation than linear regression, likely because they better accounted for closely related individuals. Although the lowest p-values from the LMM-based methods meet the conventional criterion of 5e-08 for genome wide significance (Table S7), our interpretation is that none of these variants warrant further investigation. Low observed minor allele frequencies (<2%) are inconsistent with those expected for variants responsible for pigmentation differences between the African and Native American populations because the frequencies of alleles responsible for population differences are expected to be highly differentiated between these source populations.

Additional Native American hypopigmenting alleles of significant effect size remain to be identified. Previously characterized variants do not explain this difference. It is possible that multiple hypopigmenting variants of small effect sizes are together required to reach Native American and/ or East Asian levels of hypopigmentation, individually having insufficient effect to detect in the Kalinago, given our power limitations. If this is the case, multiple variants are required to explain the observed net difference in pigmentation. Alternatively, if there are variants with large effect sizes, it appears likely that they were not genotyped and are poorly tagged by the genotyped SNPs. Additional work will be required to find hypopigmentation alleles of significant effect size that are responsible for the lighter color of Native Americans.

## Material and Methods

### Ethics Statement

The study was reviewed and approved by the Kalinago council and institutional review boards of Penn State University (29269EP), Ross University, and the Dominica Ministry of Health (H125). Informed consent was obtained from each participant enrolled in the study, and in the case of minors, consent was also obtained from a parent or guardian.

### Recruitment

Participants from among the Kalinago populations were recruited with the help of nurses from the Kalinago Territory in 2014. Recruitment took place throughout the territory’s 8 hamlets. Place and date of birth, reported ancestry of parents and grandparents, number of siblings, and response to sun exposure (tanning ability, burning susceptibility) were obtained by interview. Hair color and texture and eye color (characterized as black, brown, gray, blue, green, hazel, no pigment) were noted visually but not measured quantitatively.

### Skin Reflectometry

Skin reflectance was measured using a Datacolor CHECK_PLUS_ spectrophotometer and converted to melanin unit as we have previously described (Ang et al., 2012; Diffey et al., 1984). To minimize the confounding effects of sun exposure and body hair, skin color measurements were measured on each participant’s inner arm, and the average of triplicate measurements was generated. Before skin color measurements were taken, alcohol wipes were used to minimize the effect of dirt and/or oil. In order to minimize blanching due to occlusion of blood from the region being measured, care was taken not to apply only sufficient pressure to the skin to prevent ambient light from entering the scanned area (Fullerton et al., 1996).

### DNA Collection

Saliva samples were collected using the Oragene Saliva kit, and DNA was extracted using the prepIT.L2P kit, both from DNA Genotek (Ottawa, Canada). DNA integrity was checked by agarose gel electrophoresis and quantitated using a NanoDrop spectrophotometer (Thermo Fisher Scientific, Waltham, MA). Further quantification was done using Qubit Fluorometer (Thermo Fisher Scientific, Waltham, MA) as needed, following manufacturer instructions.

### Genotyping

Oculocutaneous albinism variants previously identified in African and Native Americans (Carrasco et al., 2009; King et al., 2003; Stevens et al., 1997; Yi et al., 2003) were amplified by PCR in all albino individuals as well as control samples using published conditions. Selected alleles of *SLC24A5, SLC45A2, OCA2 and MFSD12* were amplified in all sampled individuals as described in Table S8. Amplicons generated by 30 cycles of PCR using an Eppendorf thermocycler were sequenced (GeneWiz, South Plainfield, NJ) and the chromatograms viewed using Geneious software.

Illumina SNP genotyping using the Infinium Omni2.5-8 BeadChip was performed for all the individuals sampled. This was performed in three cohorts, using slightly different versions of the array, and the results combined. Due to ascertainment differences between the cohorts, analysis is presented here only for the combined sample. After quality control to eliminate duplicates and monomorphic variants, and to remove variants and individuals with genotype failure rates > 0.05, 358 Kalinago individuals and 1 638 140 unique autosomal SNPs remained.

### Whole exome sequencing of albino individual and obligate carrier

In order to identify the causative variant for albinism in the Kalinago, 2 samples (one albino individual and one parent) were selected for whole exome sequencing. Following shearing of input DNA (1 microgram) using a Covaris E220 Focused-ultrasonicator (Woburn, MA), exome enrichment and library preparation was done using the Agilent SureSelect V5+UTR kit (Santa Clara, CA). The samples were sequenced at 50x coverage using a HiSeq 2500 sequencer (Illumina, San Diego, CA).

The *fastq* files were aligned back to Human Reference Genome GRCh37 (HG19) using BWA(Li and Durbin, 2009) and bowtie (Langmead et al., 2009). Candidate SNP polymorphisms were identified using GATK’s UnifiedGenotyper (McKenna et al., 2010), while the IGV browser was used to examine the exons of interest for indels (Thorvaldsdottir et al., 2013). Variants with low sequence depth (< 10) in either sample were excluded from further consideration.

### Computational analysis

Basic statistics, merges with other datasets, and association analysis by linear regression were performed using plink 1.9 (Chang et al., 2015; Purcell et al., 2007). Phasing and imputation, as well as analysis of regions of homozygosity by descent and identity by descent were performed with Beagle 4.1 (Browning and Browning, 2013, 2007), using 1000 Genomes Project (1KGP) phased data (The 1000 Genomes Project Consortium, 2015) as reference.

The genotyped individuals were randomly partitioned into nine subsets of 50 or 51 individuals (n=50 subsets) in which no pair exhibited greater than second-order relationship (PI_HAT > 0.25 using the -- genome command in plink). Using the same criteria, a maximal subset of 184 individuals was also generated (n=184 subset).

Principal components analysis (PCA) was performed using the smartpca program (version 13050) in the eigensoft package (Price et al., 2006). For comparison to HGDP populations, Kalinago samples were projected onto principal components calculated for the HGDP samples alone. For use as covariates in association analyses, the n=184 subset was used to generate the PCA, and the remaining individuals were projected onto the same axes.

Admixture analysis was performed using the ADMIXTURE program (Alexander et al., 2009; Zhou et al., 2011). Each of the nine n=50 Kalinago subsets was merged with the N=940 subset of HGDP data (Li et al., 2008; Rosenberg, 2006) for analysis (349,923 SNPs) and the outputs combined, averaging ancestry proportions for the common HGDP individuals across runs. These results were used in figures. Separately, two-stage admixture analysis started with the averaged estimated allele frequencies and then employed the projection (--P) matrix outputs to estimate individual ancestry for the combined Kalinago sample. Individual ancestries estimated using both methods, as well as those estimated from a thinned subset of 50,074 SNPs were in good agreement, consistent with standard errors estimated by bootstrap analysis, although sample-wide averages differed slightly. Cross-validation is enable by adding the --cv to the ADMIXTURE command.

For association analyses we removed the three-albino individuals and excluded SNPs with minor allele frequency < 0.01. For conventional association analysis by linear regression, the standard additive genetic model included sex, the first 10 PCs, and genotypes of rs1426654 (*SLC24A5*), rs16891982 (*SLC45A2*) and the albino variant rs797044784 (*OCA2*) as covariates (Table S9). Linear mixed model (LMM) analysis was performed using the mlma module of GCTA (Yang et al., 2011) with the --mlma-no-preadj-covar flag to suppress calculation using residuals. Two Genetic Relatedness Matrix (GRM) were used: a standard GRM calculated using GCTA’s --make-grm command, and an ancestry-aware GRM calculated using relationships deduced by REAP (Thornton et al., 2012) that utilized the output of the two-stage Admixture analysis. For linear regression only, P-values were adjusted for statistic inflation by genomic control using the lambda calculated from the median chi-square statistic.

Statistical power was estimated by simulation, using a subset of genotyped SNPs. Starting with the 349,923 SNPs used for ancestry analysis, the averaged P matrix from ADMIXTURE analysis at K=4 provided an initial estimate of allele frequencies in AFR and NAM ancestral populations. 10,233 SNPs exhibited differentiation of 0.7 or greater between these populations, a value chosen as a reasonable minimum population differentiation for causative variants. After removal of SNPs for which predicted Kalinago sample frequencies deviated by more than 0.1 from observed values and those with adjusted P < 0.1, 8766 SNPs remained. Phenotypes were simulated by randomly selecting one of these SNPs and adding a defined effect size to the observed phenotype. Simulated datasets were then analyzed with plink using the standard genetic model.

Statistical analysis of pigmentary effect of albinism involved fitting parameters to an additive model for the sample containing carriers but lacking albino individuals, applying the same model to the albino individuals, and comparing residuals for the albinos and the other individuals.

Local ancestry analysis of the region containing the albinism allele was performed using the PopPhased version of rfmix (v1.5.4) with the default window size of 0.2 cM (Maples et al., 2013). A subset of 1KGP data served as reference haplotypes for European, African and East Asian populations, and the Native American ancestry segments of the admixed samples as determined by (Martin et al., 2017a) were combined to generate synthetic Native American reference haplotypes. For estimates of individual ancestry, Viterbi outputs for each window were averaged across all autosomes.

## Data Availability Statement

The whole exome sequencing and whole genome SNP genotyping data underlying this article cannot be shared publicly due to the privacy of individuals and stipulation by the Kalinago community. Only de-identified filtered SNP data used in analyses will be shared. Additional data will be shared on request to the corresponding author, pending approval from the Kalinago Council.

## Acknowledgments

We would like to thank the Kalinago Council, Dominica Ministry of Health, nurses at the Kalinago Territory, Salybia Mission Project and the Kalinago community for their assistance and participation in this study. We would also like to acknowledge faculty of Ross University, Portsmouth, Dominica (now Bridgetown, Barbados), especially Drs. Gerhard Meisenberg (retired) and Liris Benjamin of Ross University in helping us to obtain the necessary IRB approval for fieldwork. This work was supported by the Hershey Rotary Club, Jake Gittlen Laboratories for Cancer Research and Department of Pathology for funding portions of this project. We would also like to acknowledge members of the Cheng Lab for their constructive comments and input.

