## Supplement File for "Native American Ancestry and Pigmentation Allele Contributions to Skin Color in a Caribbean Population"

**Supplementary Figures**

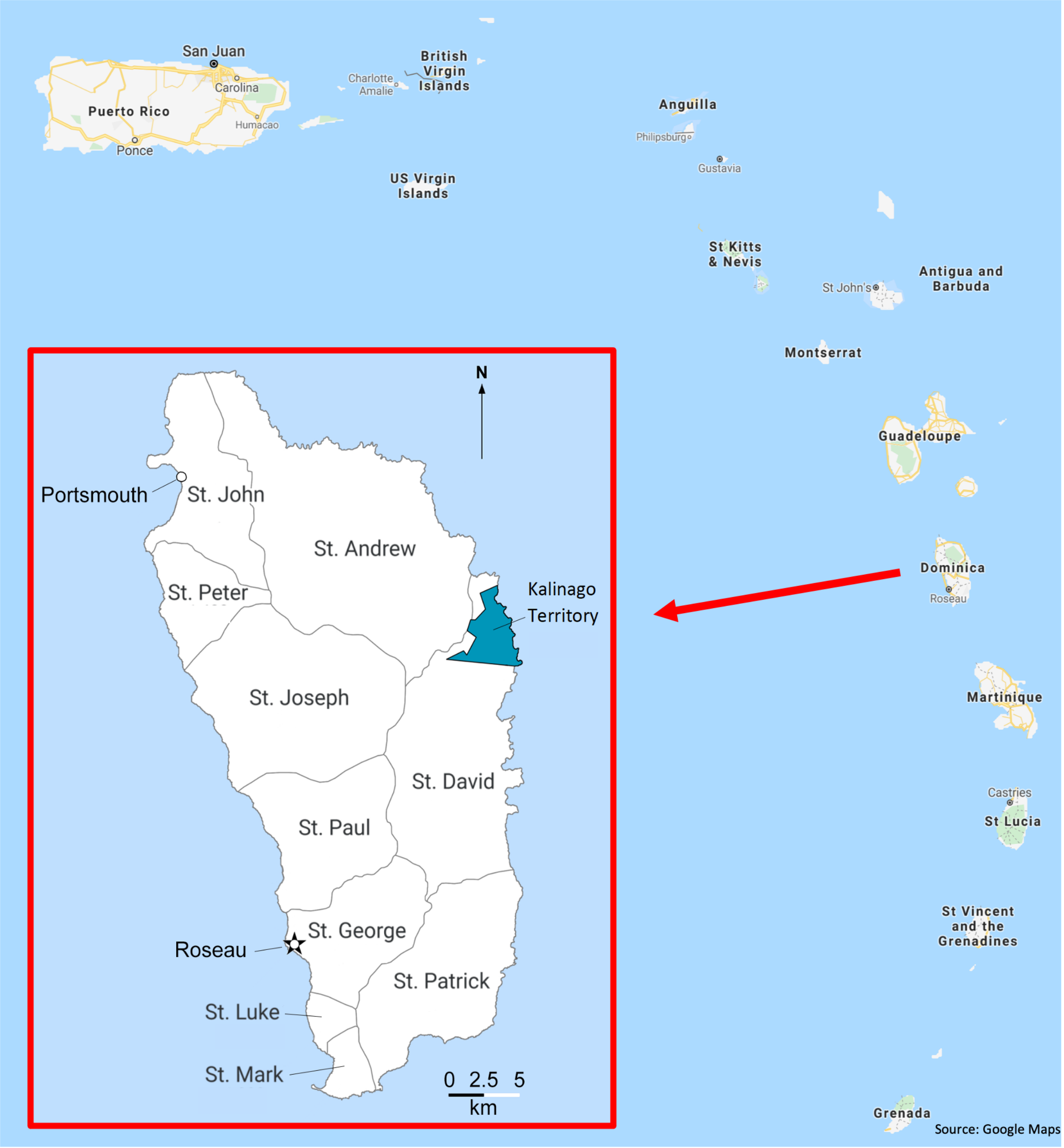


**Figure S1. Map showing the location of Kalinago Territory in the Commonwealth of Dominica**. Dominica, also known as *Wai'tu kubuli* in the Kalinago language, is clustered with the Leeward Islands in the Lesser Antilles archipelago of the Caribbean Sea. Main map situates Dominica within the Eastern Caribbean. Inset shows Dominica, with location of Kalinago Reservation (blue) in relation to parishes and principal towns. (Map modified from SESA CROP Report and Google Maps.)

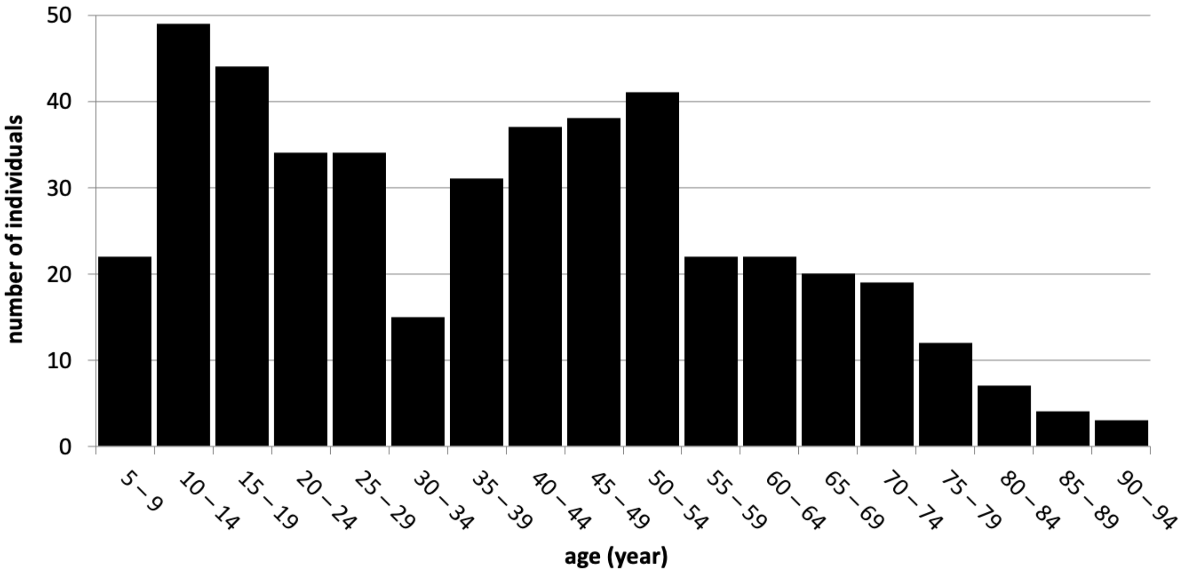


**Figure S2. Age distribution of sampled Kalinago individuals**. Histogram shows age in years at last birthday for all sampled individuals for whom this information was collected (n=455).

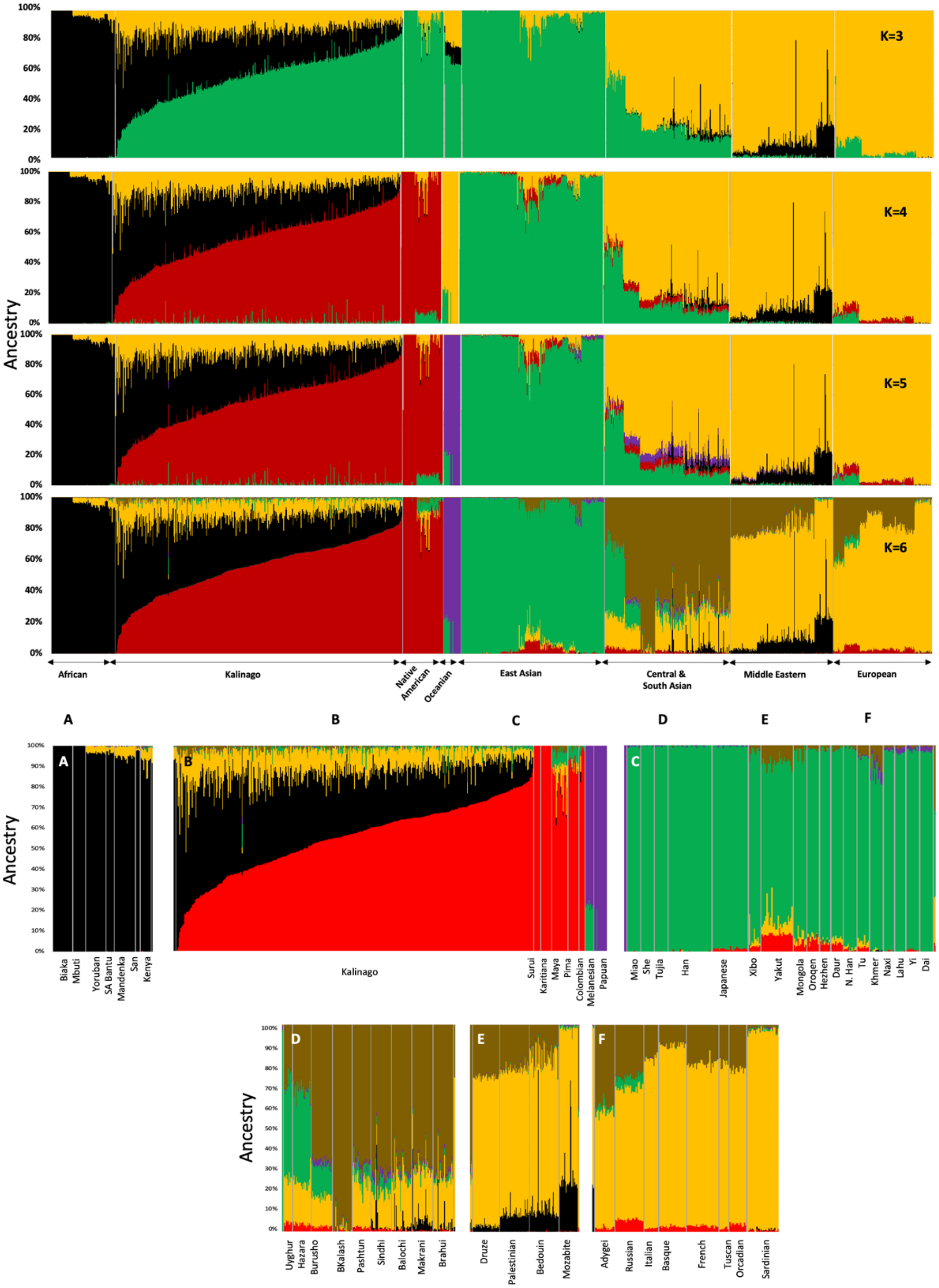


**Figure S3: Admixture plot of Kalinago compared to Human Genome Diversity Project** data from K=3 to K=6. Expended admixture plot at K=6 labeled each of the populations used, from panel A-F.

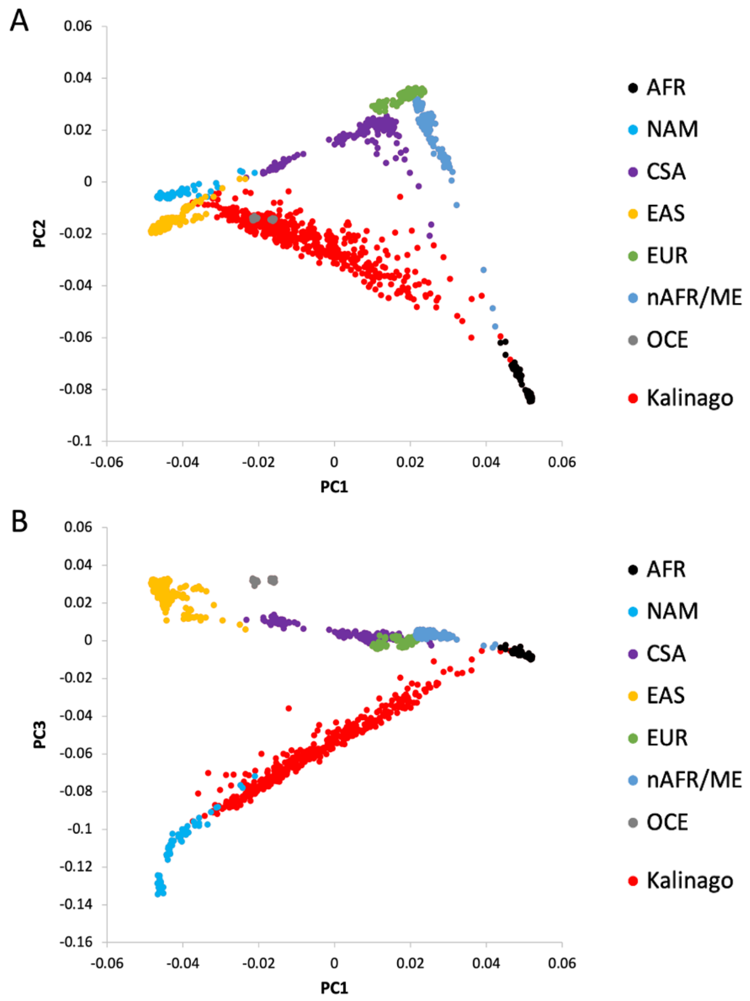


**Figure S4. Principal Components Analysis of Kalinago and comparison populations**. PCA analysis was performed on HGDP sample (940 individuals), with 458 Kalinago individuals projected on the same axes. **A**, PC1 and PC2; **B,** PC1 and PC3. In both panels, HGDP individuals are colored to indicate cluster membership (AFR, African; nAFR/ME, Northern Africa and Middle East; EUR, Europe; CSA, Central and Southern Asia; EAS, East Asia; OCE, Oceania; NAM, Native American). Ancestry was represented by the first 10 principal components because AFR and NAM ancestries are not independent of each other. The first PC correlated strongly with AFR or NAM ancestry ($r^2$ 0.94 and 0.97, respectively), but also with EUR ancestry ($r^2 = 0.32$). Several other principal components displayed considerably lower levels of correlation with ancestry ($r^2 < 0.1$ for EUR and $r^2 < 0.05$ for EAS). Individuals homozygous for the albino variant were excluded from association analyses. Association analysis did not reveal any novel variants that reached genome-wide significance, after correction for statistic inflation. The inflation factor (lambda) for the full genotyped sample excluding the albinos (n=444) sample was 1.349. Values of lambda for the nine N=50 subsets ranged from 1.001 to 1.184 (median 1.075), suggesting that the elimination of second order relatives did not remove all effects of relatedness.

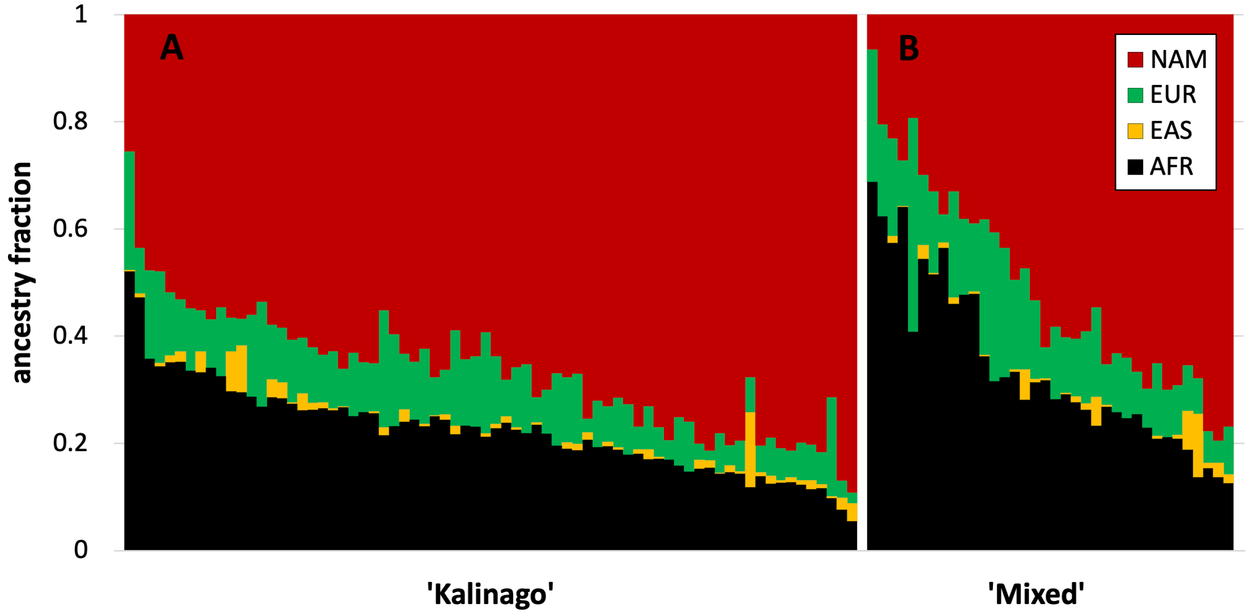


**Figure S5. Ancestry distribution as function of community-defined ancestry**. Individual ancestry fraction was estimated using Admixture (K=4) as described. Individuals identified as **A** 'Kalinago' (n=72) have higher NAM and lower AFR and EUR ancestry than those identified as **B** 'Mixed' (n=36). Despite considerable overlap in ancestry proportions between individuals, the distributions are distinctly different. Compared to individuals identified as “Mixed,” those identified as “Kalinago” have on average more Native American ancestry (67% vs 51%), less European ancestry (10% vs 14%), and less African ancestry (23% vs 34%). Similarly, the phenotypic distributions of the two groups differed.

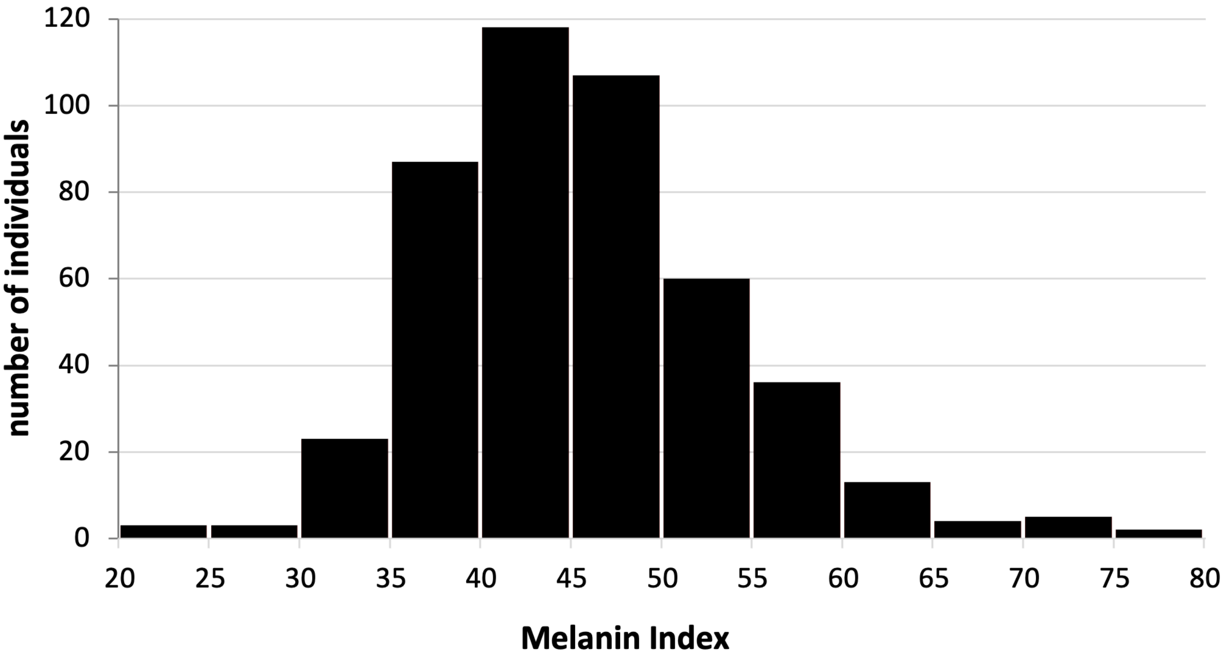


**Figure S6: Skin color distribution of the Kalinago from Commonwealth of Dominica.** We collected 462 Kalinago who live in the Kalinago Reservation. Each participant was asked a set of questions about their ancestry, gave their saliva sample, and have their skin color measured under their arm.

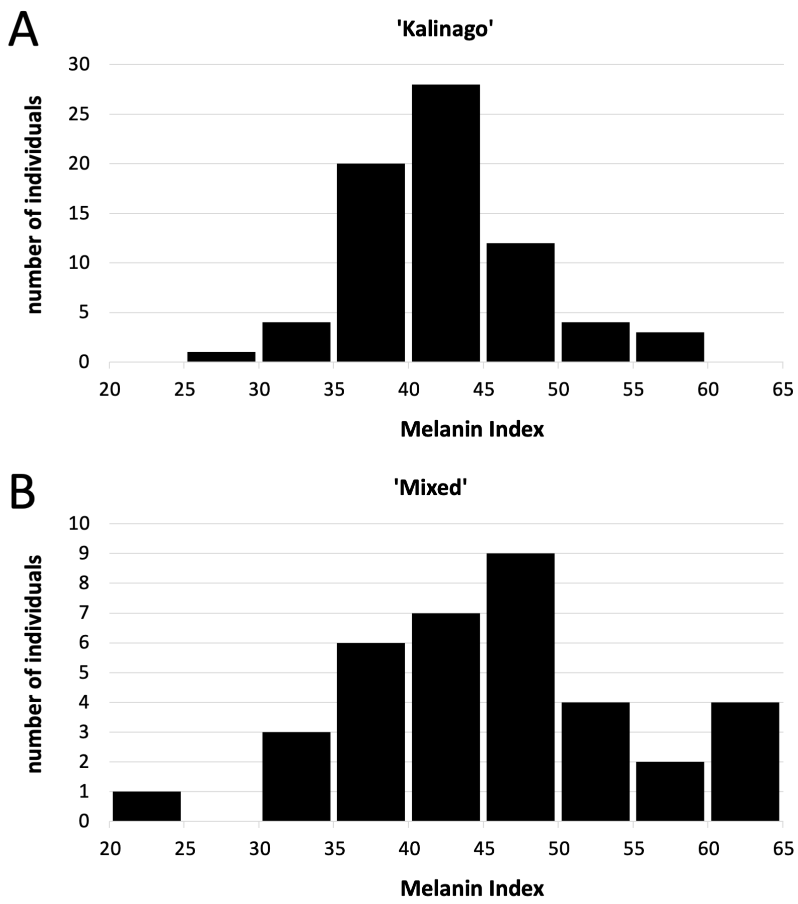


**Figure S7: Melanin Index distribution as function of community-described ancestry.** Individuals described as (**A**) “Kalinago” (n=72) were slightly lighter and had a narrower MI distribution (42.5 ± 5.6, mean ± SD) than those described as (**B)** “Mixed” (45.8 ± 9.6).

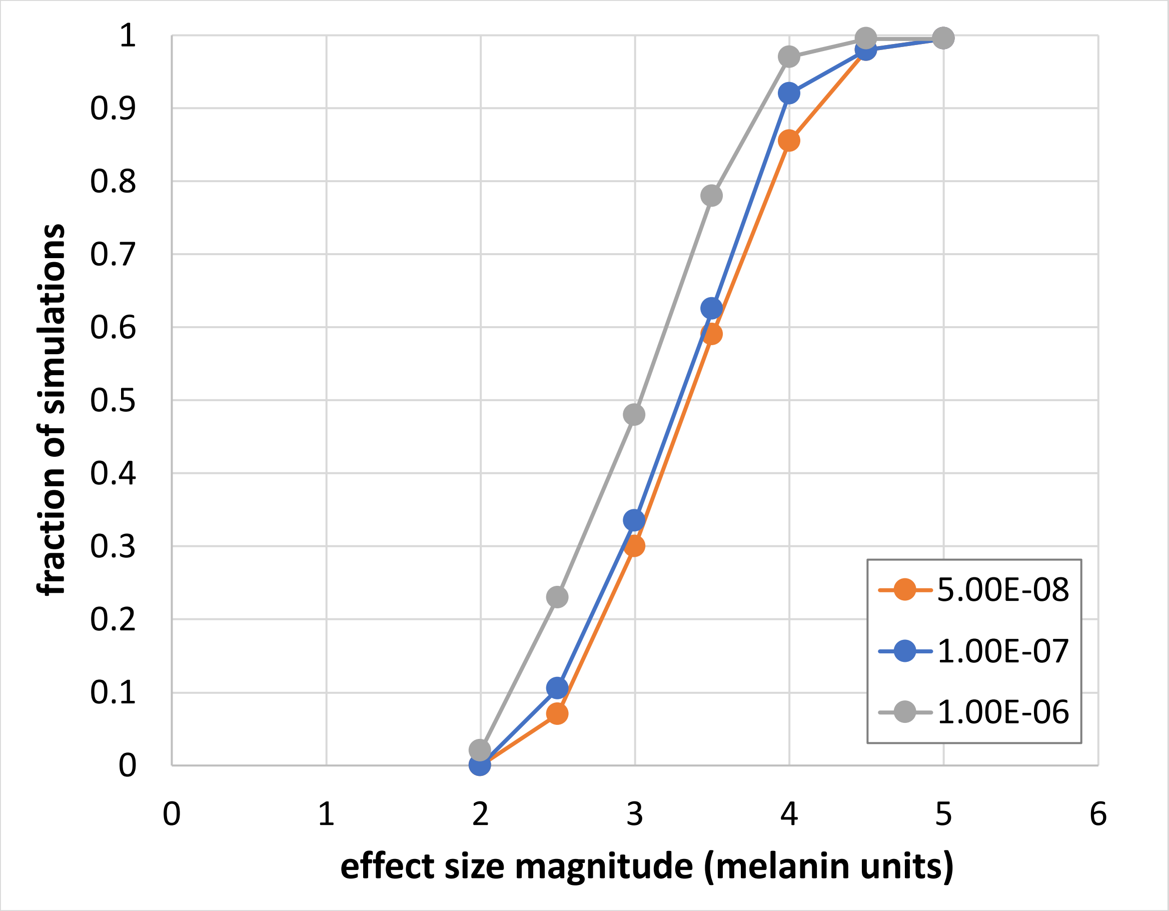


Figure S8. **Estimated power for GWAS using Kalinago sample**. Simulations were performed as described in Methods, using genotyped SNPs with estimated frequency difference between African and Native American ancestral populations of at least 0.7 and adjusted p-value of at least 0.1.

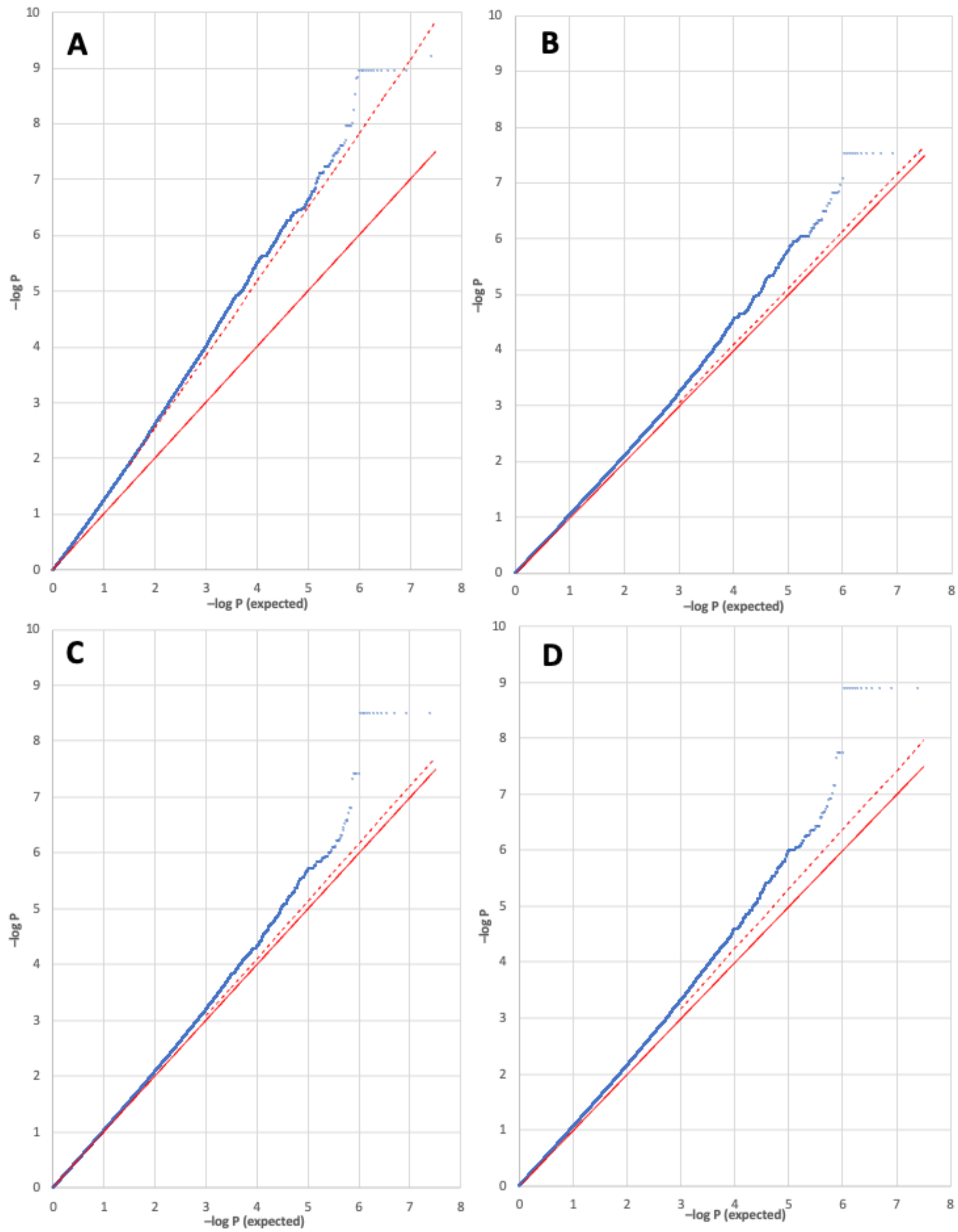


**Figure S9. Q-Q plots for association analyses.** All estimates (blue dots) were calculated using $SLC24A5^{A111T}$, $SLC45A2^{L374F}$, $OCA2^{NW273KV}$, and sex as covariates. Plotted values are not corrected for statistic inflation. Red line shows expected values; dashed red line shows expected values based on statistic inflation (lambda) calculated from median. A) Linear regression with 10 principal components (PCs) included as covariates, Lambda = 1.342; B) Linear Mixed Model (LMM) with no PCs, standard Genetics Related Matrix (GRM), Lambda = 1.024; C) LMM with 10 PCs, standard GRM, Lambda = 1.031; D) LMM with 10 PCs, REAP GRM. Lambda = 1.068.

### Supplementary Tables

**Table S1. Sample Demographics.**

| Category | Entire sample (N=461) |
|---|---|
| **Sex** | |
| male | 244 |
| female | 217 |
| **Age** | |
| range | 6 to 93 |
| mean (SD) | 39 (21.5) |
| median | 39 |
| **Paternal ancestry** | |
| reported[a] | 432 |
| named | 193 |
| sampled[b] | 49 |
| **Maternal ancestry** | |
| reported[a] | 437 |
| named | 244 |
| sampled[c] | 128 |

[a] community-described ancestry collected.

[b,c] values from reported genealogy; 75 fathers and 146 mothers as determined by genotyping.

**Table S2: Summary of Kalinago ancestry from admixture analysis (n=458).** NAM = Native American, AFR = African, EUR = European, CSA = Central & South Asian, EAS = East Asian, OCE = Oceanian. At K=3, NAM, EAS, and OCE are not distinguishable.

<table>
<tr><th>K-value</th><th>AFR</th><th>NAM</th><th>EAS</th><th>OCE</th><th>EUR</th><th>CSA</th></tr>
<tr><td>3</td><td>0.304</td><td colspan="3">0.552</td><td colspan="2">0.144</td></tr>
<tr><td>4</td><td>0.318</td><td>0.549</td><td colspan="2">0.011</td><td colspan="2">0.122</td></tr>
<tr><td>5</td><td>0.318</td><td>0.548</td><td>0.011</td><td>0.002</td><td colspan="2">0.121</td></tr>
<tr><td>6</td><td>0.318</td><td>0.548</td><td>0.012</td><td>0.002</td><td>0.110</td><td>0.010</td></tr>
</table>

Table S3. Ancestry proportions estimated using different approaches.

| estimation approach | AMR | AFR | EUR | EAS |
|---|---|---|---|---|
| Admixture (subsets, K=4) | 0.549 | 0.318 | 0.122 | 0.011 |
| Admixture (two stage, K=4) | 0.541 | 0.316 | 0.126 | 0.016 |
| rfmix (4 clusters) | 0.553 | 0.313 | 0.125 | 0.009 |
| rfmix (3 clusters) | 0.557 | 0.326 | 0.117 | --- |

**Table S4A. Summary by locus of albinism candidates identified through exome sequencing.** Candidates are homozygous derived in one albino and heterozygous in one obligate carrier. No nonsense, frameshift, or splice variants was detected. Our initial attempt to identify the albinism variant in the Kalinago involved targeted genotyping of the albino individuals for 28 mutations previously observed[38,39,53,54] in African or Native American albinos; these included the 2.7 kb exon 7 deletion in *OCA2* found at high frequency in some African populations. No mutation was detected using this approach.

| OCA gene | Chromosome | Variants | Missense |
|---|---|---|---|
| *OCA1 (TYR)* | 11 | 0 | |
| *OCA2* | 15 | 5 | 2 |
| *OCA3 (TYRP1)* | 9 | 0 | |
| *OCA4 (SLC45A2)* | 5 | 0 | |
| *OCA5* | 4 | 6 | 0 |
| *OCA6 (SLC24A5)* | 15 | 0 | |
| *OCA7 (LRMDA)* | 10 | 1 | 0 |

**Table S4B. Characteristics of individual candidates identified through exome sequencing.**

| Chr | rsID | Ref | Alt | f(AFR)[a] | Gene | Location/ Effect |
|---|---|---|---|---|---|---|
| 4 | rs3733437 | T | C | 0.126 | *EMCN* | intron |
| 4 | rs6826912 | T | G | 0.327 | *PPP3CA* | 3'UTR |
| 4 | rs463373 | T | C | 0.986 | *SLC39A8* | 3'UTR |
| 4 | rs439757 | C | A | 0.986 | *SLC39A8* | 3'UTR |
| 4 | rs223495 | A | G | 0.399 | *MANBA* | intron |
| 4 | rs3733632 | A | G | 0.819 | *TACR3* | 5'UTR |
| 10 | rs7911113 | A | G | 0.476 | *LRMDA* | intron |
| 15 | rs1800419 | A | G | 0.629 | *OCA2* | synonymous |
| 15 | rs1800401 | G | A | 0.126 | *OCA2* | *R305W* |
| 15 | rs797044784[b] | CCAG | GACC | 0.002 | *OCA2* | *NW273KV* |
| 15 | rs73375883 | G | A | 0.203 | *OCA2* | intron |
| 15 | rs972334 | G | A | 0.217 | *OCA2* | intron |

[a] Overall frequency for non-reference allele in seven 1KGP African populations.
[b] 1KGP describes this variant as four consecutive SNPs rs549973474, rs569395077, rs538385900 and rs558126113.

**Table S8. Amplification conditions used for genotyping Kalinago samples for the selected alleles**.

| Gene & Variant | Primer Sequence | PCR Annealing Temperature (°C) |
|---|---|---|
| *SLC24A5*$^{A111T}$<br>rs1426654 | Fwd- CTCACCTACAAGCCCTCTGC<br>Rev- AATTGCAGATCCAAGGATGG | 55 |
| *SLC45A2*$^{L374F}$<br>rs16891982 | Fwd- CCTGCTGGGACTCATCCATC<br>Rev- AGCAGAGTGCATGAGAAGGG | 55 |
| *OCA2*$^{NW273KV}$<br>rs797044784 | Fwd- AGAGTCCCAGATGGTGTCTCA<br>Rev- AGGTCAGACTCCTTTAAACG | 53 |
| *OCA2*$^{R305W}$<br>rs1800401 | Fwd- AGAGGGAGGTCCCCTAACTG<br>Rev- ATCTCAAGCCTCCCTGACTG | 53 |
| *MFSD12*$^{Y182H}$<br>rs2240751 | Fwd- CCCAGGTGGAATAGCAGTGAG<br>Rev- AGTGGTTGGAATCACCTGTCA | 61 |
